# Relationships between acute race-induced changes in creatine kinase activity and blood plasma myoglobin concentration and race performance in mountain bike and road cyclists

**DOI:** 10.1101/2021.05.17.444439

**Authors:** Paulina Hebisz, Jacek Borkowski, Rafał Hebisz

## Abstract

The aim of this study was to determine if the changes in plasma creatine kinase (CK) activity and myoglobin (MB) concentrations as markers of muscle damage differ between competitive road (*n* = 14) and mountain bike (*n* = 11) cyclists and if these biochemical markers show a relationship with real-world race performance. CK and MB were measured from blood samples collected 2 hours before race start and 1 hour after race completion and the change in pre- and post-race difference was calculated (ΔCK and ΔMB). An incremental exercise test was used to determine maximal oxygen uptake, maximal aerobic power, and power output at the second ventilatory threshold. Post-race CK and MB increased in the whole group of cyclists. Although the magnitude of change in CK was similar in both road and mountain bike cyclists, only the increase in road cyclists was significant. MB significantly increased only in mountain bike cyclists. Multiple regression analysis revealed a significant association between both road and mountain bike race performance and ΔCK and ΔMB. The other significant predictors for mountain bike race performance were maximal aerobic power (W·kg^-1^) and power output at the second ventilatory threshold (W·kg^-1^) and for road race performance both maximal oxygen uptake (l·min^-1^) and power output at the second ventilatory threshold (W). In conclusion, mountain bike racing was associated with an increase in MB whereas road racing with an increase in CK, with the post-race changes in CK and MB related to race performance as high ΔCK and low ΔMB were obtained by better-performing cyclists.

## Introduction

The mountain bike (MTB) cycling in the Olympic format (XCO) is a highly-intensive race lasting approximately 1 hour 30 min and is performed on technically demanding courses. During competition, average heart rate exceeds 90% maximal heart rate [1,2] and power output measured during uphill climbs is frequently above maximal aerobic power [1,3]. Biochemical analysis during simulated MTB racing showed that blood pH decreases to 7.3 and blood lactate concentration increases to 7.5 mmol/l [4]. Road cycling is of significantly longer duration and performed at a lower intensity. Average power output at the elite level is approximately 200 W [5,6] with average heart rate in the range of 50–60% maximal heart rate depending on the terrain [7].

Aside from differences in duration and exercise intensity, competitive road and MTB cycling also differ in muscle activity patterns. Existing literature has noted that the difficult terrain conditions of MTB can elicit significant eccentric muscle contractions for shock attenuation and handling particularly during downhill sections [2-4] and may compound energy expenditure even when little to no pedaling is required [4]. This type of varied and prolonged muscle action including maximal bouts during uphill climbs can severely stress skeletal muscle [8], and cause structural damage within the muscle fiber [9], which had previously been confirmed in MTB cyclists [10]. The varied static, eccentric and concentric phases of MTB [4] can be compared with the principally concentric muscle action that characterizes road cycling [11].

The differences in road cycling and MTB racing formats (distance covered, terrain, and uphill/downhill phases) can also be reflected in pacing strategy, in which road cyclists show more linear and uniform pacing and power output [12] than MTB cyclists [1]. Accordingly, the disparities in the physiological demands and activity profiles of these two competitive cycling modalities suggest that the skeletal muscle damage should occur slower but may develop in a longer race time in road cycling relative to MTB. One of the causes of muscle damage could be cumulative oxidative stress [13]. Recent studies have used biochemical analysis to provide a more composite assessment of post-race oxidative damage [13]. It would seem prudent to continue this line of research to further assess and quantify post-race muscle damage in order to elucidate the physiological profiles of competitive road cycling and MTB. Research has found creatine kinase (CK) activity and myoglobin (MB) concentration to serve as useful markers of acute muscle damage and be correlated with training status [8]. These biomarkers of muscle status are particularly attractive as previous studies have suggested that they can also be used to predict cycling performance [10].

Understanding the relationship between potential changes in post-race CK activity and MB concentration and real-world race performance can have several practical implications. Previous studies have demonstrated a strong relationship between winning performance in road cycling and MTB and maximal oxygen uptake, power output at lactate threshold, and gross mechanical efficiency [14,15]. These variables reflect the aerobic energy potential of athletes and resistance to fatigue [9]. However, the onset of fatigue is modulated by a number of factors outside those which are cardiovascular, related to energy supply, or biomechanical, with current research highlighting the role of muscle trauma particularly during prolonged, high-intensity exercise [9]. A greater understanding of the relationship between the biochemical markers of post-race muscle damage and race performance could improve knowledge on the physiological determinants of success in MTB and road cycling.

The aim of this investigation was to therefore determine if the changes in CK activity and MB concentration differ between competitive road and MTB cyclists and if these biochemical markers show a relationship with race performance. It was hypothesized that the specificity of MTB racing would result in greater CK activity and MB concentration and that the elevated levels of these fatigue-indicating biochemical markers would be negatively associated with race performance.

## Materials and methods

### Participants

The study involved well-trained competitive road (*n* = 14) and MTB (*n* = 11) cyclists. All participants were included in the same racing age category: under 23 (U23). No differences in baseline anthropometric measurements were observed between the two groups (Table 1).

**Table 1.** Anthropometric characteristics of the mountain bike and road cyclists.

|  | <u>Mountain bike cyclists</u> |  |  | <u>Road cyclists</u> |  |  | Cohen's *d* |
| --- | --- | --- | --- | --- | --- | --- | --- |
|  | Mean ± *SD* | CI 95% |  | Mean ± *SD* | CI 95% |  |  |
|  |  | Upper | Lower |  | Upper | Lower |  |
| **Age [y]** | 19.1 ± 1.3 | 20.0 | 18.3 | 19.4 ±1.6 | 20.4 | 18.5 | 0.21 |
| **BM [kg]** | 68.1 ± 5.5 | 71.8 | 64.4 | 71.1 ± 4.8 | 73.8 | 68.3 | 0.58 |
| **BH [m]** | 1.79 ± 0.04 | 1.81 | 1.76 | 1.77 ± 0.04 | 1.80 | 1.75 | 0.50 |
| **BMI** | 21.3 ± 1.2 | 22.2 | 20.5 | 22.6 ± 1.79 | 23.6 | 21.6 | 0.85 |

BM – body mass; BH – body height; BMI – body mass index; SD – standard deviation; CI – confidence interval;

The study design was approved by the institutional review board and conducted in accordance with the ethical standards established by the Declaration of Helsinki. Written informed consent was obtained from all participants and their guardians after the study details, procedures, and benefits and risks were explained.

### Design and Procedures

Race data were collected during a National Cup event in which the cyclists competed in their respective discipline. Two days after race completion the aerobic capacity and performance of the cyclists was evaluated in controlled laboratory conditions at an exercise laboratory (PN-EN ISO 9001:2001 certified).

#### Race characteristics

The MTB cyclists competed in cross-country olympic (XCO) race format with the start and finish located at 320 m above sea level. Six laps were completed on a 4.8 km course with 180 m of elevation. The road cyclists competed in a one-day mass start race with the start and finish at 210 m above sea level. Course distance was 17.5 km with 160 m of elevation and lapped eight times. The cyclists were instructed to provide a maximal effort with emphasis on obtaining a high finish. They were familiarized with the course one day before competition by performing three (MTB) or two (road) practice laps. During the familiarization session, the participants were instructed to perform at an intensity below the second ventilatory threshold (VT2). The day preceding the familiarization session the cyclists rested and refrained from any exercise. Upon race completion, the official results (finish times) were posted and the difference (T_D_) between the winning time and each participant’s finish time was calculated as a measure of performance.

#### CK and MB determination

On the day of competition, a 0.5 ml sample of arterialized capillary blood from the fingertip was collected 2 hours before race start and 1 hour after race completion to determine CK activity and MB concentration. The blood samples were immediately centrifuged at 1000g for 15 min at 4°C for plasma recovery. These samples were stored in Eppendorf tubes at -30°C and analyzed once all samples were collected. CK activity was determined with an EC 2.7.3.2 assay kit (Biosystems, Spain). MB concentration was determined using the Human Myoglobin Matched Antibody Pair Kit AB215407 (Abcam, UK) with the included protein standard. ExtrAvidin-peroxidase conjugate (Sigma-Aldrich) diluted to 1:1000 was added to the plate and incubated for 1 hour at room temperature. The plate was then washed in phosphate-buffered saline containing 0.05% Tween 20. Afterwards, 0.4 mg/ml o-phenylenediamine and 0.3% H_2_O_2_ (v/v) dissolved in 0.1 M citrate buffer (pH 5.0) were added. The enzyme reaction was stopped after 30 min by adding 100 μl of a 1 M solution of H_2_SO_4_ and the absorbance was read at 490 nm (as the primary wavelength) using an Epoch TM spectrophotometer and integrated software (Bio-Tek Instruments, USA). The difference in pre- and post-race measures of CK and MB were then calculated (ΔCK and ΔMB, respectively). Creatine kinase activity and myoglobin concentration measured 1 hour after the race were corrected for plasma volume changes. For this purpose, pre- and post-race hematocrit (HCT) value and hemoglobin (HGB) concentrations were also determined prior to centrifugation with an ABX Micros OT 16 Analyser (Horiba, Poland). These measures were used to assess the percent change in blood plasma volume (%ΔPV) in accordance with Dill and Costill [16]:

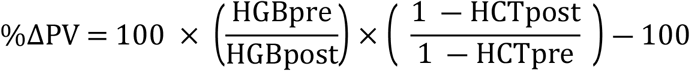

#### Incremental exercise test

An incremental exercise test (IXT) was administered on an Excalibur Sport cycle ergometer (Lode B.V., Netherlands) two days after race completion. The device was calibrated before each trial and the test began at a starting workload of 50 W that was increased every 3 min by 50 W until volitional exhaustion. If the participant was unable to complete a 3-min stage, 0.28 W was subtracted for each missing second from the obtained level of power to determine absolute and relative (per kg of body mass) maximal aerobic power (Pmax). Respiratory function was measured during the test with a face mask connected to a Quark gas analyzer (Cosmed, Italy). Calibration was performed prior to each trial with a reference gas mixture of carbon dioxide (5%), oxygen (16%), and nitrogen (79%). Tidal air was analyzed on a breath-by-breath basis to determine oxygen uptake (VO_2_), maximal oxygen uptake (VO_2_max), carbon dioxide excretion (VCO_2_), and minute pulmonary ventilation (VE). These measures were averaged over 30-s intervals. Absolute and relative VO_2_max were calculated based on VE and the composition of expired air. The second ventilatory threshold (VT2) was defined by V-slope analysis of VO_2_ and VCO_2_ [17] and absolute and relative power output at VT2 was determined.

### Statistical Analysis

The Statistica 13.1 software package (Statsoft, USA) was used to analyze the data. The distribution of the data set was screened for normality using the Shapiro–Wilk test and means ± standard deviations were calculated for each variable. Student’s *t* test for independent samples was used to identify the between-group differences for the anthropometric and IXT-obtained measures. Differences in CK and MB between the groups and time points were tested using repeated-measures analysis of variance (ANOVA) with Duncan’s test post hoc when appropriate. Significance was accepted for all procedures at *p* < 0.05 and upper and lower confidence intervals (95% CI) are presented. The relationship between T_D_ and the biochemical and IXT-obtained variables was determined using multiple regression analysis. The explanatory variables were randomly included to develop the most optimal model. Only the regression models with the best fit and accepted at a significance level of *p* < 0.05 are presented.

## Results

Obtained in incremental exercise test absolute and relative VO_2_max, absolute Pmax, and absolute and relative power output at VT2 were significantly greater whereas T_D_ was significantly lower in the group of road cyclists compared with the mountain bike cyclists (Table 2). There was a significant main effect of time for CK activity (*F* = 15.6, *p* = 0.001, *η*^*2*^ = 0.40) and MB concentration (*F* = 17.2, *p* = 0.000, *η*^*2*^ = 0.43). A significant group × time interaction was observed for MB concentration (*F* = 4.5, *p* = 0.046, *η*^*2*^ = 0.16). The results of Duncan’s test show a significant post-race increase in CK activity in the road cyclists and a post-race increase in MB concentration in the MTB cyclists (Table 3). Three regression models were selected that best explain the relative contribution of the different independent variables to the difference in winning time and finish time (Table 4).

**Table 2.** Incremental exercise test and race characteristics of the mountain bike and road cyclists.

|  | <u>Mountain bike cyclists</u> |  |  | <u>Road cyclists</u> |  |  |  |
| --- | --- | --- | --- | --- | --- | --- | --- |
| | Mean ± *SD* | CI 95% | | Mean ± *SD* | CI 95% | | Cohen's |
|  |  | Upper | Lower |  | Upper | Lower | *d* |
| **VO<sub>2</sub>max**<br>[l·min<sup>-1</sup>] | 4.28 ± 0.51 | 4.62 | 3.94 | 4.82 ±0.33* | 5.01 | 4.63 | 1.26 |
| **VO<sub>2</sub>max**<br>[ml·min<sup>-1</sup>·kg<sup>-1</sup>] | 62.9 ± 5.2 | 66.4 | 59.4 | 67.9 ± 4.4* | 70.4 | 65.4 | 1.04 |
| **Pmax** [W] | 378.2 ± 41.2 | 405.9 | 350.5 | 415 ± 27.5* | 430.9 | 399.1 | 1.05 |
| **Pmax**[W·kg<sup>-1</sup>] | 5.56 ± 0.52 | 5.91 | 5.21 | 5.86 ± 0.51 | 6.16 | 5.56 | 0.58 |
| **VT2** [W] | 264.5 ± 43.6 | 235.2 | 293.8 | 315.4±40.5* | 338.8 | 291.9 | 1.21 |
| **VT2** [W·kg<sup>-1</sup>] | 3.89 ± 0.55 | 4.26 | 3.52 | 4.46 ±0.67* | 4.85 | 4.07 | 0.93 |
| **T<sub>D</sub>** [s] | 644.3 ± 383.6 | 902.0 | 386.5 | 204.9 ±255.9* | 352.6 | 57.1 | 1.35 |

VO<sub>2</sub>max – maximal oxygen uptake; Pmax – maximal aerobic power; VT2 – power output at the second ventilatory threshold; T<sub>D</sub> – difference between winning time and finish time; SD – standard deviation; CI – confidence interval; \* – between-group difference significant at p<0.05

**Table 3.** Pre- and post-race CK activity and MB concentration and blood plasma volume.

|  | <u>Pre-race</u> |  |  | <u>Post-race</u> |  |  | Cohen's |
| --- | --- | --- | --- | --- | --- | --- | --- |
| | Mean ± *SD* | CI 95% | | Mean ± *SD* | CI 95% | | *d* |
|  |  | Upper | Lower |  | Upper | Lower |  |
| Mountain bike cyclists |  |  |  |  |  |  |  |
| **CK** [u·l<sup>-1</sup>] | 320.1 ±182.0 | 442.4 | 197.8 | 392.3 ± 237.3 | 551.7 | 179.6 | 0.43 |
| **MB** [ng·ml<sup>-1</sup>] | 21.0 ± 14.3 | 30.6 | 11.4 | 31.6* ± 20.1 | 45.1 | 18.1 | 1.69 |
| **ΔPV** [%] | | | | -4.1 ± 9.3 | 2.2 | -10.3 | |
| Road cyclists |  |  |  |  |  |  |  |
| **CK** [u·l<sup>-1</sup>] | 267.1 ±131.7 | 343.2 | 191.1 | 336.9* ± 143.7 | 419.8 | 253.9 | 0.51 |
| **MB** [ng·ml<sup>-1</sup>] | 17.8 ±4.3 | 20.3 | 15.4 | 21.3 ±3.9 | 23.6 | 19.0 | 0.85 |
| **ΔPV** [%] | | | | 0.1 ± 13.9 | 8.1 | -7.9 | |

CK – plasma creatine kinase activity; MB – plasma myoglobin concentration; ΔPV – change in plasma volume; SD – standard deviation; CI – confidence interval; \* – pre- and post-race difference significant at p<0.05

**Table 4.** Partial correlation coefficients between the race time difference and independent variables among the three best regression models.

|  | VO<sub>2</sub>max | VO<sub>2</sub>max | ΔCK | ΔMB | Pmax | Pmax | VT2 | VT2 |
| --- | --- | --- | --- | --- | --- | --- | --- | --- |
|  | [l·min<sup>-1</sup>] | [ml·min<sup>-1</sup>·kg<sup>-1</sup>] | [u·l<sup>-1</sup>] | [ng·ml<sup>-1</sup>] | [W·kg<sup>-1</sup>] | [W] | [W] | [W·kg<sup>-1</sup>] |
| T<sub>1</sub> | --- | --- | -0.65 | --- | -0.62 | --- | --- | -0.56 |
| T<sub>2</sub> | --- | -0.51 | -0.88* | 0.59 | --- | --- | --- | --- |
| T<sub>3</sub> | -0.42 | --- | -0.47 | 0.77* | --- | --- | -0.40 | --- |

T<sub>1</sub> – time difference between MTB winning time and MTB cyclist's finish time; T<sub>2</sub> – time difference between MTB winning time and MTB cyclist's finish time after excluding the two best MTB cyclists; T<sub>3</sub> – time difference between the road race winning time and road cyclist's finish time; VO<sub>2</sub>max – maximal oxygen uptake; ΔCK – pre- and post-race difference in creatine kinase activity; ΔMB – pre- and post-race difference in myoglobin concentration; Pmax – maximal aerobic power; VT2 – power output at the second ventilatory threshold; \* – correlation significant at $p < 0.05$

Model 1 (*R* = 0.93, *R* ^*2*^= 0.87, *F* = 16.01, *p* = 0.002, *SEE* = 163.5) – explains the time difference (T_1_) between MTB winning time and MTB cyclist’s finish time with the regression equation:

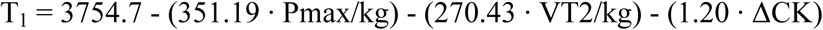

Model 2 (*R* = 0.91, *R*^*2*^ = 0.84, *F* = 7.72, *p* = 0.02, *SEE* = 177.1) – explains the time difference (T_2_) between MTB winning time and MTB cyclist’s finish time after excluding the two best cyclists (described in further detail in Discussion) with the regression equation:

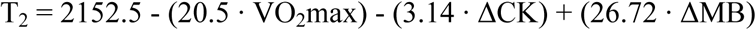

Model 3 (*R* = 0.87, *R*^*2*^ = 0.75, *F* = 6.80, *p* = 0.008, *SEE* = 160.2) – explains the time difference (T_3_) between the road race winning time and road cyclist’s finish time with the regression equation:

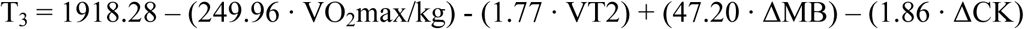

## Discussion

The results showed a significant increase in CK only in the road cycling group. It is worth noting that the magnitude of change in CK was similar in both the road and MTB groups with similar effect sizes (Cohen’s *d*) and that no significant between-group differences were found at either pre- or post-race. In turn, MB increased significantly only in the MTB group, suggesting more pronounced fatigue after MTB racing compared with road racing. The lack of congruency in the direction of changes in CK activity and MB concentration may be conditioned by a variety of factors that should be mentioned.

Although a commonly used marker of exercise-induced muscle damage, CK activity is frequently assessed several hours after exercise in contrast with the sample collection time in the present study [18]. One reason is due to the difference in the molecular masses of CK and MB (82 kDa and 17.5 kDa, respectively), in which the greater mass of CK prolongs post-exercise clearance from muscle tissue to blood [19]. For example, in a study involving trained endurance runners, peak blood CK activity were observed 28 ± 11 hours after starting a 100 km race [20]. Other studies have reported peak blood CK activity 8 hours after a single strength training session with no changes in time-to-peak level after 4 months of continual training [21]. In the present study, blood samples were collected no later than 1 hour after completion due to the logistical considerations of the participants, in which many had to immediately depart after concluding the race. This early collection of post-race CK may not reflect actual post-exercise fatigue status as circulating CK may be still low. Another consideration is that CK composition is two-fold greater in fast-twitch compared with slow-twitch muscle fiber [22,23]. Conversely, MB concentration in slow-twitch fibers is 1.5:1 times greater than in fast-twitch fibers [24,25]. As a result, changes in CK levels may more accurately reflect fatigue status in fast-twitch muscle (which could have been similar in both the road and MTB groups) whereas the change in MB concentration may provide a better composite picture of slow-twitch fatigue (which was greater in the MTB group).

Regression analysis showed that ΔCK is a variable negatively related to the time difference between winning time and the participant’s finish time in each of the models; cyclists with greatest ΔCK obtained the lowest T_D_. The increase in CK activity after exercise is ordinarily attributed to muscle cell membrane and tissue damage due to large muscle tension and eccentric exercise [8,18]. The intense ATP hydrolysis with a concurrent decrease in intracellular pH can lead to dysfunction of the Ca^2+^-ATPase pump, causing a large influx of calcium ions that can increase proteolytic enzyme activity. Such conditions can degrade cell membrane proteins, increasing cell membrane permeability and promoting an outflux of CK into the blood [18]. It is known that fast-twitch fibers are better adjusted to ATP synthesis via the glycolytic pathway than slow-twitch fibers [26]. It is therefore possible that ΔCK variability in the present group of cyclists is dependent on the recruitment of fast-twitch fibers for ATP production to support slow-twitch fibers.

When comparing the regression models, the relationship between ΔCK and T_D_ is stronger in the model for MTB cyclists than road cyclists. This could partly be attributed to the specificity of MTB racing as it involves significant intensive eccentric exercise [2] that can result in significant muscle cell damage particularly to Z-band muscle fibers [8,18]. By excluding the two best MTB cyclists, an even stronger relationship between ΔCK and the time difference between winning time and finish time was found for MTB cyclists. This finding is best interpreted within the context of the race, in which these two highest-ranked MTB cyclists (smallest T_D_) completed the first half of the race in the leading group composed of 4 cyclists. As a result, by sharing the time with the fastest riders at least in the first phase of the race, these cyclists maintained a pace that may have been at a lower intensity than their abilities. This suggests that the increase in CK activity could be lower among better-performing cyclists, which in these two cyclists ΔCK was 31.5 u/l and 0.0 u/l.

Interestingly, the regression coefficient of ΔMB was positive (in contrast with ΔCK) and an independent variable only in the models predicting the difference between winning time and participant’s finish time in the road race and in the MTB race only after excluding the two best participants. This suggests that it is race-induced damage to slow-twitch fibers which negatively affects winning time (increasing T_D_). It is also noteworthy that the two best MTB cyclists mentioned above showed ΔMB values (16.1 ng/ml and 40.2 ng/ml) at a magnitude similar to the slowest cyclists. This may suggest greater involvement of slow-twitch fibers during maximal exercise which minimizes the recruitment of fast-twitch fibers and indicative that the onset of fatigue may be more closely related to disruption in slow-twitch muscle. Another potential explanation for this may be attributed to muscle fiber type transformation in the best-performing cyclists, in which fast-twitch fibers transition to show properties of slow-twitch fibers [27].

Finally, the developed regression models confirm the relative contribution of VO_2_max, Pmax, and power output at VT2 to performance in road and MTB cycling [6,14,15]. While the presence of these variables in the generated models is not surprising, the lack of significant partial correlation with performance is in contradiction with previously reported findings involving cyclists. Borszcz et al. [28] observed a high correlation between mean power output in individual time trials and Pmax and VO_2_max. Heuberger et al. [29] analyzed the predictive value of power at the lactate threshold with real-world road race performance and found moderate to strong correlations depending on the method used to calculate the lactate threshold. However, a study by Impellizzeri et al. [30] found MTB race performance to be moderately correlated only with power output at VT2 normalized to body mass and not VO_2_max. Our findings suggest that ΔCK and ΔMB can be used to ascertain training status and race performance in road and MTB cyclists and complement typical physiological variables such as Pmax, VO_2_max, and power output at VT2.

## Conclusions

The main findings of the study demonstrated that MTB racing was associated with an increase in MB concentration whereas road racing with an increase in CK activity. The post-race changes in CK and MB appear to be related to performance (race time) as high ΔCK and low ΔMB were obtained by better-performing cyclists (smallest time difference between winning time and finish time). However, this association was not found when considering the two highest-ranked MTB cyclists who had the best times but lowest ΔCK.

## Acknowledgments

Our immense gratitude to Michael Antkowiak for his translation of the manuscript and language assistance.

## Author Contributions

**Conceptualization:** Paulina Hebisz, Jacek Borkowski, Rafal Hebisz

**Data curation:** Rafał Hebisz, Jacek Borkowski

**Formal analysis:** Rafal Hebisz

**Funding acquisition:** Paulina Hebisz, Rafal Hebisz

**Investigation:** Paulina Hebisz, Jacek Borkowski, Rafal Hebisz

**Methodology:** Paulina Hebisz, Jacek Borkowski, Rafal Hebisz

**Project administration:** Paulina Hebisz

**Resources:** Paulina Hebisz

**Software:** Rafal Hebisz

**Supervision:** Paulina Hebisz

**Validation:** Jacek Borkowski

**Visualization:** Paulina Hebisz, Jacek Borkowski, Rafal Hebisz

**Writing – original draft:** Paulina Hebisz, Jacek Borkowski, Rafal Hebisz

**Writing – review & editing:** Paulina Hebisz, Jacek Borkowski, Rafal Hebisz

## Notes

### Competing Interest Statement

The authors have declared no competing interest.

## References

1. Granier C, Abbiss CR, Aubry A, Vauchez Y, Dorel S, Hausswirth C, et al. Power output and pacing during international cross-country mountain bike cycling. Int J Sports Physiol Perform. 2018;24:1–22.

2. Impellizzeri FM, Marcora S. The physiology of mountain biking. Sports Med. 2007;37(1):59–71.

3. Macdermid PW, Stannard S. Mechanical work and physiological responses to simulated cross country mountain bike racing. J Sports Sci. 2012;30(14):1491–501.

4. Hays A, Devys S, Bertin D, Marquet LA, Brisswalter J. Understanding the physiological requirements of the mountain bike cross-country olympic race format. Front Physiol. 2018;9:1062.

5. Ebert TR, Martin DT, Stephens B, Withers RT. Power output during a professional men’s road-cycling tour. Int J Sports Physiol Perform. 2006;1:324–35.

6. Mujika I, Padilla S. Physiological and performance characteristics of male professional road cyclists. Sports Med. 2001;31(7):479–87.

7. Padilla S, Mujika I, Orbañanos J, Santisteban J, Angulo F, José Goiriena J. Exercise intensity and load during mass-start stage races in professional road cycling. Med Sci Sports Exerc. 2001;33(5):796–802.

8. Brancaccio P, Lippi G, Maffulli N. Biochemical markers of muscular damage. Clin Chem Lab Med. 2010;48(6):757–67.

9. Abbiss CR, Laursen PB. Models to explain fatigue during prolonged endurance cycling. Sports Med. 2002;35(10):865–98.

10. Hebisz R, Hebisz P, Borkowski J, Zatoń M. Effects of concomitant high-intensity interval training and sprint interval training on exercise capacity and response to exercise-induced muscle damage in mountain bike cyclists with different training backgrounds. Isokinet Exerc Sci. 2019;27(1):21–9.

11. Bijker KE, de Groot G, Hollander AP. Differences in leg muscle activity during running and cycling in humans. Eur J Appl Physiol. 2002;87(6):556–61.

12. Vogt S, Heinrich L, Schumacher YO, Blum A, Roecker K, Dickhuth HH, et al. Power output during stage racing in professional road cycling. Med Sci Sports Exerc. 2006;38(1):147–51.

13. Córdova A, Sureda A, Albina ML, Linares V, Bellés M, Sánchez DJ. Oxidative stress markers after a race in professional cyclists. Int J Sport Nutr Exerc Metab. 2015;25(2):171–8.

14. Lee H, Martin DT, Anson JM, Grundy D, Hahn AG. Physiological characteristics of successful mountain bikers and professional road cyclists. J Sports Sci. 2002;20(12):1001–8.

15. Wilber RL, Zawadzki KM, Kearney JT, Shannon MP, Disalvo D. Physiological profiles of elite off-road and road cyclists. Med Sci Sports Exerc. 1997;29(8):1090–4.

16. Dill DB, Costill DL. Calculation of percentage changes in volumes of blood, plasma, and red cells in dehydration. J Appl Physiol. 1974;37(2):247–8.

17. Beaver WL, Wasserman K, Whipp BJ. A new method for detecting anaerobic threshold by gas exchange. J Appl Physiol. 1986;60(6):2020–7.

18. Baird MF, Graham SM, Baker JS, Bickerstaff GF. Creatine-kinase- and exercise-related muscle damage implications for muscle performance and recovery. J Nutr Metab. 2012;2012:960363.

19. Perryman MB, Strauss AW, Buettner TL, Roberts R. Molecular heterogeneity of creatine kinase isoenzymes. Biochim Biophys Acta. 1983;747(3):284–90.

20. Stäubli M, Roessler B, Köchli HP, Peheim E, Straub PW. Creatine kinase and creatine kinase MB in endurance runners and in patients with myocardial infarction. Eur J Appl Physiol Occup Physiol. 1985;54(1):40–5.

21. Hurley BF, Redmond RA, Pratley RE, Treuth MS, Rogers MA, Goldberg AP. Effects of strength training on muscle hypertrophy and muscle cell disruption in older men. Int J Sports Med. 1995;16(6):378–84.

22. Tai PW, Fisher-Aylor KI, Himeda CL, Smith CL, Mackenzie AP, Helterline DL, et al. Differentiation and fiber type-specific activity of a muscle creatine kinase intronic enhancer. Skelet Muscle. 2011;1:25.

23. Yamashita K, Yoshioka T. Profiles of creatine kinase isoenzyme compositions in single muscle fibres of different types. J Muscle Res Cell Motil. 1991;12:37–44.

24. Jansson E, Sylvén C. Myoglobin concentration in single type I and type II muscle fibres in man. Histochemistry. 1983;78(1):121–4.

25. Khan MA. Histochemical characteristics of vertebrate striated muscle: a review. Prog Histochem Cytochem. 1976;8(4):1–48

26. Karlsson J, Sjödin B, Jacobs I, Kaiser P. Relevance of muscle fibre type to fatigue in short intense and prolonged exercise in man. Ciba Found Symp. 1981;82:59–74.

27. Putman CT, Xu X, Gillies E, MacLean IM, Bell GJ. Effects of strength, endurance and combined training on myosin heavy chain content and fibre-type distribution in humans. Eur J Appl Physiol. 2004;92(4-5):376–84.

28. Borszcz FK, Tramontin AF, de Souza KM, Carminatti LJ, Costa VP. Physiological correlations with short, medium, and long cycling time-trial performance. Res Q Exerc Sport. 2018;89(1):120–5.

29. Heuberger JA, Gal P, Stuurman FE, de Muinck Keizer WA, Mejia Miranda Y, Cohen AF. Repeatability and predictive value of lactate threshold concepts in endurance sports. PLoS One. 2018;13(11):e0206846.

30. Impellizzeri FM, Marcora SM, Rampinini E, Mognoni P, Sassi A. Correlations between physiological variables and performance in high level cross country off road cyclists. Br J Sports Med. 2005;39(10):747–51.

